# Super-resolved spatial transcriptomics reveals early hippocampal RNA localization changes in a mouse model of Alzheimer’s disease

**DOI:** 10.1101/2025.09.24.678295

**Authors:** Yaara Karasik, Hadar Eger Ronsky, Tal Goldberg, Yeela Seri Huri, Nofar Schottlender, Tom Ben-Mor, Modi Safra, Moshe Shenhav, Hila Zak, Michal Danino-Levi, Yaeli Rosenberg, Alex Glick, Noam Feldman, Noa Slater, Irit Gottfried, Uri Ashery, Shahar Alon

**Affiliations:** Faculty of Engineering, Bar-Ilan University, Israel; Institute for Nanotechnology and Advanced Materials (BINA), Bar-Ilan University, Israel; The Multidisciplinary Brain Research Center, Bar-Ilan University, Israel; School of Neurobiology, Biochemistry and Biophysics, The George S. Wise Faculty of Life Sciences, Tel Aviv University, Israel; Sagol School of Neuroscience, Tel Aviv University, Israel

## Abstract

Cell-type-specific changes in gene expression and RNA localization are hallmarks of Alzheimer’s disease (AD) and other neurodegenerative disorders, yet spatial dysregulation in early disease stages remains poorly defined. Here, we applied Expansion Sequencing (ExSeq) to map the spatial distribution of 101 genes at super-resolution in the hippocampus of 4-week-old 5xFAD and wild-type (WT) mice, prior to overt pathology. We uncovered early alterations in RNA spatial organization and gene expression, including 23 genes showing altered localization without changes in abundance in the 5xFAD hippocampus. Using spatial expression analysis and single-cell neighborhood analysis, we identified cell-type- and region-specific molecular programs associated with synaptic function, neuroinflammation, and metabolic stress that differed between 5xFAD and WT mice. Spatial RNA velocity further revealed state differences influenced by local cell to cell interactions. Together, these results suggest that RNA positioning and transcriptional programs are perturbed at early disease stages. Finally, we provide the full super-resolution ExSeq dataset as an open resource for spatial and cell-type-specific analyses in early Alzheimer’s disease research.

**Highlights:**

- Super-resolved transcriptomic profiling of the hippocampus at early disease stages
- Identification of 23 genes with altered spatial localization without changes in abundance
- Early alterations in single-cell neighborhood organization in the 5xFAD hippocampus
- Spatial RNA velocity reveals cell-type-specific cell state differences shaped by cell-cell proximity

**Graphical abstract:** 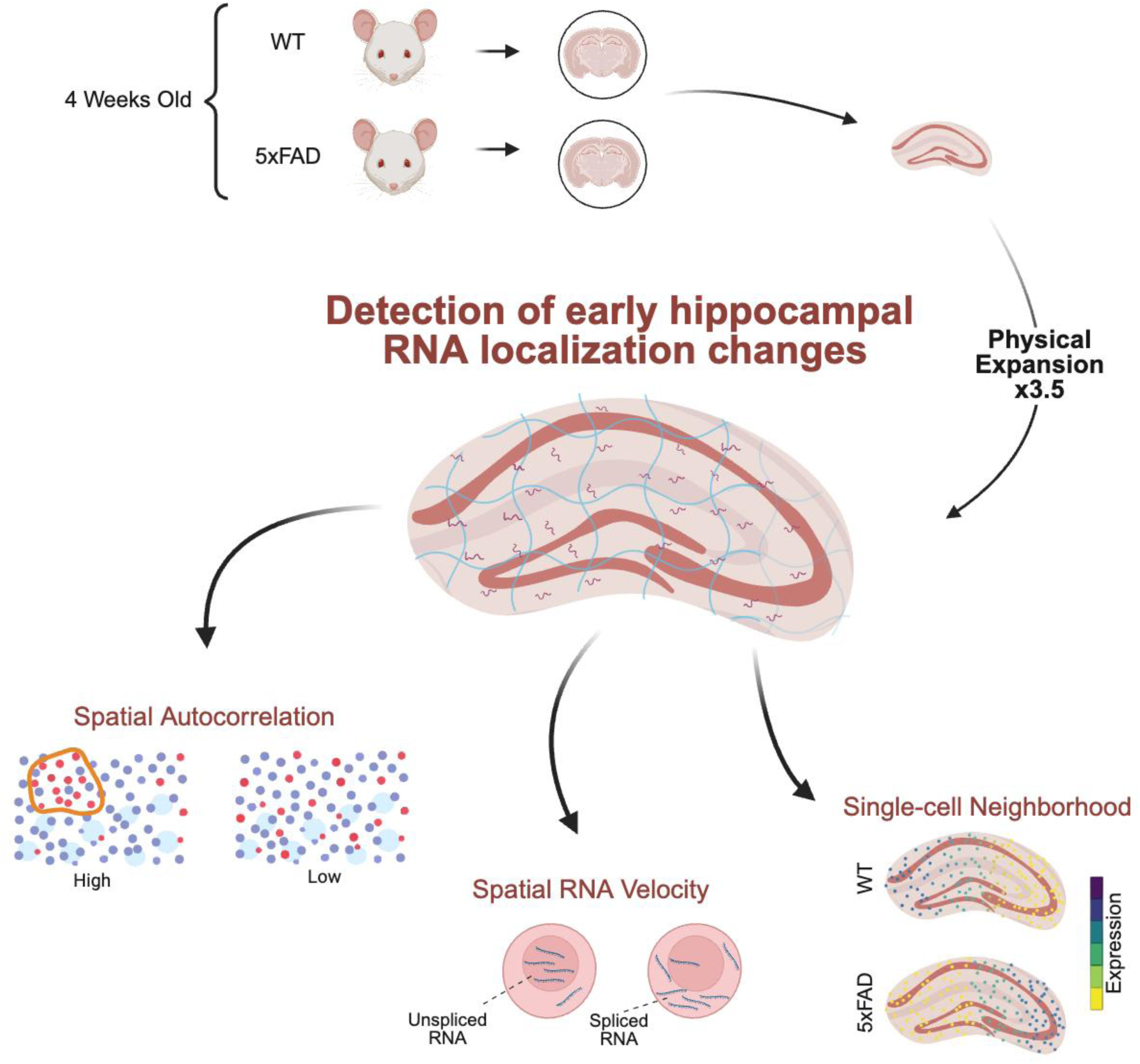

## Introduction

Alzheimer’s disease (AD) is marked by progressive synaptic and neuronal loss, with molecular pathologies such as amyloid-beta (Aβ) plaques and tau tangles^1,2^. However, molecular changes in early AD remain elusive^3–5^, particularly those that are spatially localized yet not evident from bulk expression profiles. RNA localization is tightly regulated in neurons and is essential for synaptic function and plasticity^6–9^. Mislocalization of RNA, independent of expression, may contribute to early pathogenesis^10^. Recent developments in spatial transcriptomics allow gene expression to be measured within intact tissue architecture^11^, but commercial technologies often lack single-cell resolution^12–15^.

Spatial analyses of AD molecular deficiencies using several AD mouse models: the 5xFAD^16,17^(familial AD), APP knock-in^18^, and Trem2 mutant^19^ mouse models, have largely focused on late disease stages, when pathology is already evident. For example, Visium-based^12^ profiling at 3 months and later mapped glial activation^20^; studies in APP knock-in models at ∼18 months charted plaque-proximal programs^21^; and more recently^22^, MERFISH^23,24^ atlases enabled single-cell analyses that revealed glial and neuronal transcriptomic alterations, again emphasizing late stages of the disease. To our knowledge, spatial transcriptomic analyses have not yet characterized early molecular deficiencies in AD models, particularly not with single-cell resolution. Early molecular changes have instead been described using bulk genomics: a microarray study reported hundreds of differentially expressed hippocampal and cortical genes in 4-week-old mice, progressing to an immune-dominant signature by ∼4 months^25^. Extending this effort, hippocampal RNA-seq combined with proteomics at 1, 2, and 4 months identified early, sex-biased gene dysregulation^26^.

Expansion Sequencing (ExSeq), which combines tissue expansion with in situ sequencing, enables nanoscale RNA localization and high gene throughput in thick tissue sections^27^. Here, we used ExSeq to profile the hippocampus in 4-week-old 5xFAD mice, a model of familial AD^28–30^, and age- and sex-matched wild-type littermate controls, to detect early spatial transcriptomic changes before plaque deposition.

## Results

### Framework for detecting early molecular differences in the hippocampus of an AD mouse model

We set out to identify early molecular differences in the hippocampus of 5xFAD mice compared with age- and sex-matched wild-type (WT) littermate mice, prior to overt pathology. To this end, we applied Expansion Sequencing (ExSeq), which enables quantification of both mRNA expression and spatial localization, together with three complementary spatial analysis strategies: spatial autocorrelation, single-cell neighborhood analysis, and spatial RNA velocity (Fig. 1A). At one month of age, hippocampal morphology appeared preserved and no pathology-associated markers were detected, as confocal imaging and quantification of synaptic and astrocytic markers revealed no clear differences between 5xFAD and WT mice (Fig. 1B-C, Fig. S1). At this stage no plaque accumulation was detected (Fig. S1), yet 4 weeks later, plaque load is detected in the same hippocampal areas^28,30^. It is possible that molecular changes already start at that asymptomatic stage, highlighting the need for refined spatial transcriptomic approaches such as ExSeq to uncover early, subtle molecular alterations.

**Fig. 1.**
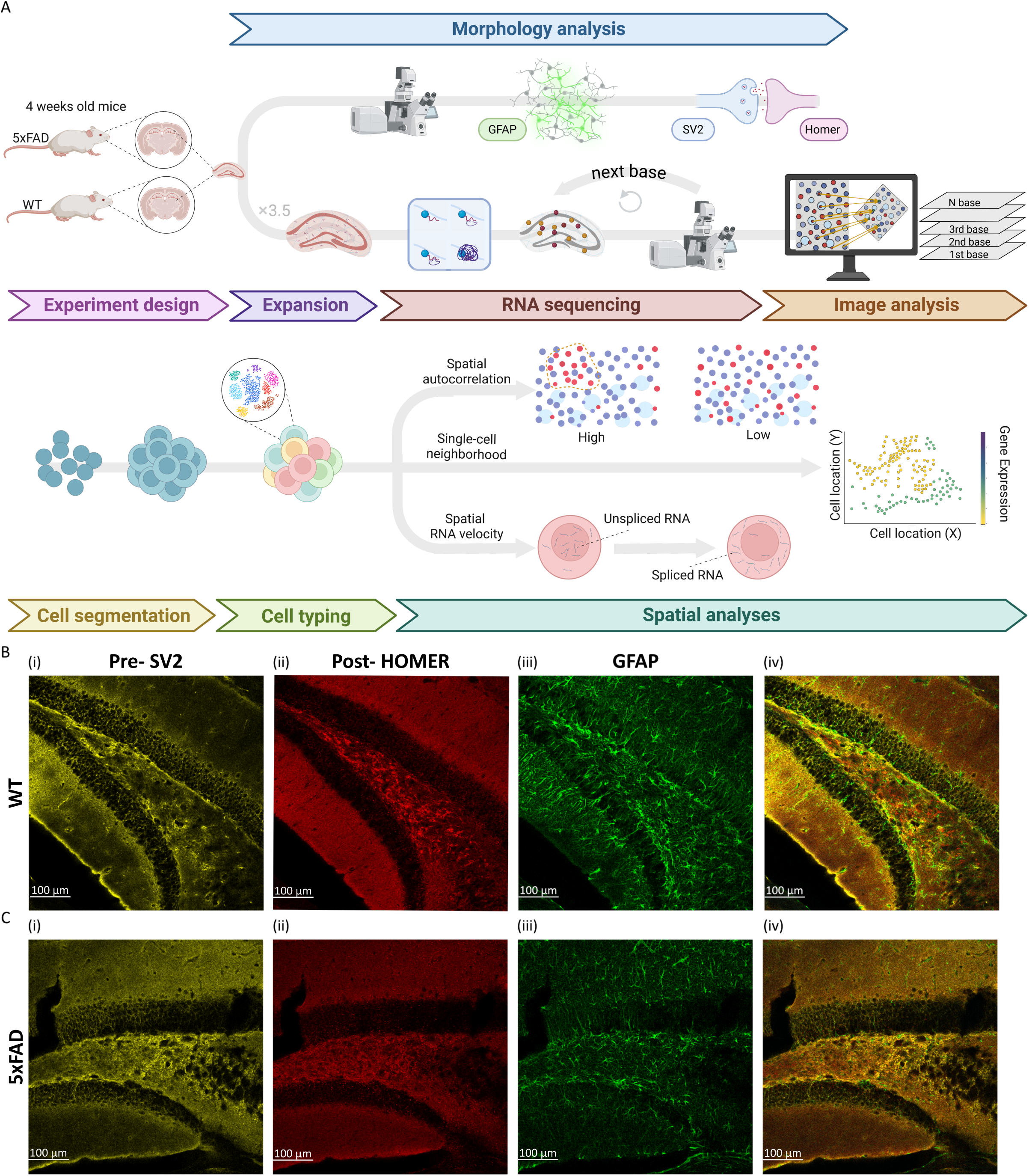
Analysis pipeline for detecting early differences between 5xFAD and WT, with hippocampal morphology preserved at this stage. **(A)** Schematic of the overall pipeline to detect early gene markers. Hippocampal sections from 5xFAD and control mice were analyzed by morphology analysis as well as super-resolved in situ sequencing. Three spatial analysis strategies: spatial autocorrelation, single-cell neighborhood analysis, and spatial RNA velocity, were applied to identify early localization changes in Alzheimer’s disease. **(B-C)** Hippocampal morphology is preserved in 1-month-old 5xFAD compared to control mice. **(B)** Representative confocal images of the hippocampus from one-month-old female WT mice at ×20 magnification. Sections were stained for the presynaptic marker SV2 (i, yellow), the postsynaptic marker Homer (ii, red), the astrocytic marker GFAP (iii, green), and (iv) merged images. **(C)** same as (B) for 5xFAD. Overall, quantification of pre- and postsynaptic markers, the astrocytic marker, and plaque load did not reveal clear differences between WT and 5xFAD at one month of age (Fig. S1).

### Super-resolved spatial sequencing of the early 5xFAD hippocampus

We profiled hippocampal sections from three female 5xFAD mice and three female wild-type (WT) mice at 4 weeks of age (n = 3 biological replicates per genotype), targeting 101 genes in 3D using ExSeq (Fig. 2A-B and Table S1). For each genotype, one mouse was processed in duplicate as technical replicates, yielding 8 total samples (4 per genotype) and 4.45 million in situ sequencing reads overall (per-sample range 122,813-962,369; group means WT: 574,498, 5xFAD: 538,956; Table S2).

**Fig. 2.**
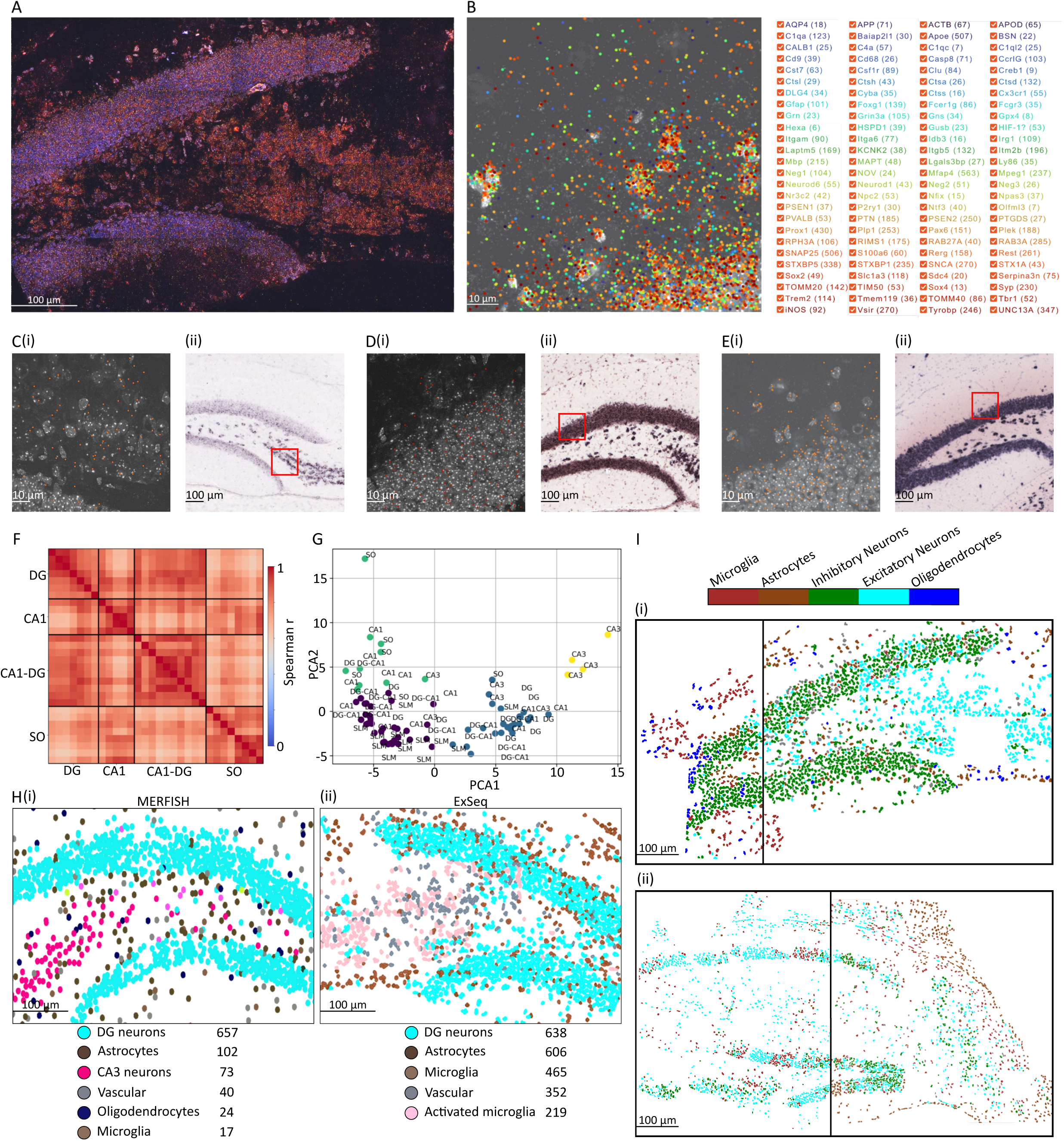
Spatial sequencing and cell type identification in brain regions relevant to early stages of AD in 5xFAD and control mice. **(A)** In situ sequencing of 101 genes in the hippocampus of a one-month-old 5xFAD mouse. A 30-µm thick hippocampal section was sequenced using ExSeq. Shown is one sequencing round with cell nuclei (DAPI, white) and two-color Illumina sequencing chemistry (red and magenta). **(B)** One representative field of view (FOV; 100 × 100 µm; all units in this study are pre-expansion) showing cell nuclei (DAPI, white) and all sequenced genes (color-coded as indicated in the legend, with overall expression in brackets). Displayed is a single Z-plane with 0.4-µm axial resolution. On average, 11,362 RNA molecules were sequenced per FOV. **(C)(i)** Example of the physical locations of one gene, *Rerg*, in a single FOV (orange dots). (C)(ii) *Rerg* expression in the dentate gyrus (purple) from a FISH experiment (Allen Brain Atlas) performed on a different biological sample. **(D)** similar to C, for *Tomm20*. **(E)** similar to C, for *Rab3a*. **(F-G)** Spatial relationships of gene expression profiles across FOVs from a 5xFAD hippocampus. **(F)** Spearman rank correlation matrix of gene expression profiles across FOVs, ordered by hippocampal region. Warmer colors (red) indicate higher similarity. Abbreviations: SO, stratum oriens; DG, dentate gyrus; CA1, cornu ammonis 1; CA1-DG, the projection dense region between the cell bodies of CA1 and DG. **(G)** Principal component analysis (PCA) of gene expression profiles followed by k-means clustering. Each FOV (dot) is colored by its assigned cluster and labeled according to its hippocampal region of origin. As expected, FOVs within the same hippocampal region group together in PCA space. Abbreviations: SLM, stratum lacunosum-moleculare; CA3, cornu ammonis 3. **(H)** Comparison of cell typing obtained with MERFISH (i) and ExSeq (ii) in similar hippocampal regions demonstrates high concordance. **(I)** Representative hippocampal regions from one-month-old control mice (i) and 5xFAD (ii), illustrating cell type identification from sequencing data and demonstrating technical reproducibility. Black vertical lines mark two independently processed samples that nevertheless produced highly similar results.

### Validation of gene localization and cell segmentation

We developed a custom ExSeq graphical user interface that enabled visualization of gene positions in 3D, and the full super-resolution dataset is provided as an open resource (Video S1). ExSeq gene positions were validated against Allen Brain Atlas FISH data (Fig. 2C-E and Fig. S2). Cell segmentation, performed using our 3D segmentation tool and correlated with single-cell reference datasets, enabled spatial identification of hippocampal cell types (Methods, Fig. S3, Table S2-S6). In situ sequenced fields of view (FOVs) from the same hippocampal region consistently clustered together in PCA space (Fig. 2F-G), providing technical validation of ExSeq profiles. Moreover, ExSeq-derived cell type distributions closely matched MERFISH atlases^23,24^, confirming accurate regional profiling (Fig. 2H). Finally, two samples independently processed from the same tissue section yielded highly similar results in both 5xFAD and WT mice (*P* = 0.02; permutation analysis, Methods), underscoring technical robustness (Fig. 2I). These validation steps established the ExSeq data and its graphical user interface as a unique tool to study gene alteration in 5xFAD mouse brain sections.

### Genes with changes in spatial distribution or expression levels

To detect alterations in spatial distribution, we computed spatial autocorrelation using Moran’s I, and assessed significance by permutation testing under the same setup (Fig. 3 and Methods). Each hippocampal region (Fig. 3A) was analyzed separately within samples, with FOVs serving as technical replicates. Differential expression analysis across hippocampal regions (Fig. 3Avi) identified tens of significant genes between 5xFAD and control, including 51 in cornu ammonis 1 (CA1), 42 in cornu ammonis 3 (CA3), 29 in the dentate gyrus (DG), 73 in stratum lacunosum-moleculare (SLM), 62 in the projection-dense CA1-DG region, 48 in stratum oriens (SO), and 5 in the hilus (Table S7). Importantly, three of these genes overlapped with those detected in a previous bulk sequencing study of 4-week-old 5xFAD versus control hippocampal tissue, with the same directionality: *C4a* and *Itgam* were upregulated in 5xFAD, while *Sox4* was downregulated^26^. Spatial autocorrelation analysis detected smaller sets of spatially modified genes (CA1 = 7, CA3 = 3, DG = 2, SLM = 25, DG-CA1 = 22, SO = 5, Hilus = 4; Table S8, and sensitivity analysis in Table S9). Most spatially modified genes overlapped with those showing expression changes, but twenty-three were exclusively spatially altered (Fig. 3A and Fig. S4): *Cst7, Fcgr3,* and *Mapt* in CA1; *Itm2b* in CA3; *C1qc, Creb1, Cyba, Fcgr3, Hif1a,* and *Stx1a* in SLM; *Acod1, Apod, Hexa, Psen1, Sox4,* and *Tomm20* in DG-CA1; *Baiap2l1, Mfap4,* and *Snap25* in SO; and *Creb1, Ctss, Fcgr3,* and *Nfix* in the Hilus. Several of these genes map onto well-defined functional axes. *Snap25* and *Stx1a* point to possible alterations in the presynaptic release machinery, consistent with disrupted SNARE-dependent vesicle fusion and synaptic integrity^31–33^. *C1qc, Cst7,* and *Fcgr3* map to the complement-microglia pathway, consistent with roles in synaptic tagging, lysosomal remodeling, and immune receptor signaling^34–38^. This analysis also identified genes involved in cellular stress responses, including *Hif1a*, a regulator of oxygen-sensitive transcription^39^; *Apod* (Fig. 3B,E), an antioxidant apolipoprotein^40^; and *Tomm20* (Fig. 3C,E), a mitochondrial import receptor^41,42^. Together, these spatially altered genes point to potential region-specific disruptions in synaptic release, immune surveillance, and metabolic resilience that were not captured by expression differences alone.

**Fig. 3.**
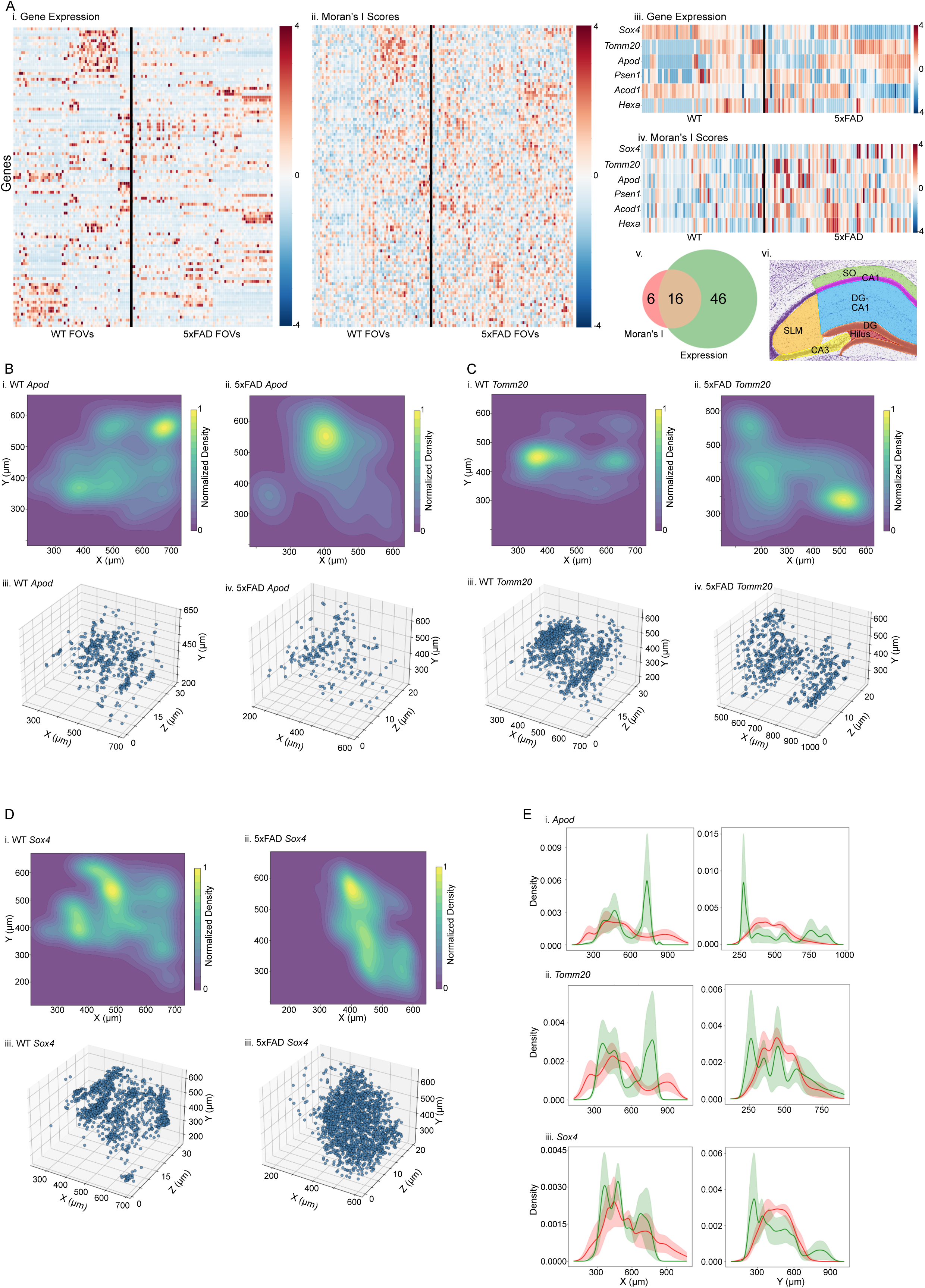
Differential spatial patterns and gene expression between 5xFAD and control hippocampus. **(A)** Spatial autocorrelation and gene expression in the DG-CA1 region. (i) Z-scaled heatmap of expression for 101 genes across 5xFAD (n=73) and control (n=54) hippocampal FOVs. (ii) Z-scaled heatmap of grid-based Moran’s I values for the same genes. (iii-iv) Heatmaps of genes significantly different between 5xFAD and control in grid-based Moran’s I but not in overall gene expression (permutation test, *q* < 0.05). (v) Venn diagram showing overlap of significant genes in the DG-CA1 region. (vi) Color-coded schematic of hippocampal subregions, including SLM, SO, CA1, CA3, DG, and Hilus. The DG-CA1 region (blue) is the region analyzed in this figure. **(B)** Spatial distribution of *Apod* in the DG-CA1 region. (i-ii) Kernel density estimation (KDE) plots of *Apod* in one representative 5xFAD sample and one control sample. (iii-iv) Scatter plots of *Apod* in the same samples. **(C)** Spatial distribution of *Tomm20*, shown as in (B). **(D)** Spatial distribution of *Sox4*, shown as in (B). **(E)** Mean density distributions of *Apod*, *Tomm20*, and *Sox4* in the DG-CA1 region along hippocampal axes in all 5xFAD (red) and control (green) samples. Solid lines show the mean KDE across samples; shaded regions indicate ± SEM. In all cases, density distributions differed significantly between 5xFAD and control (Kolmogorov-Smirnov test, *q* < 1 × 10^−9^).

### Single-cell neighborhood analysis between WT and 5xFAD

We next performed single-cell neighborhood analysis to identify genes exhibiting spatially regulated expression within individual cell types across WT and 5xFAD samples (Fig. 4A). Genes detected as spatially variable were clustered based on co-occurrence across individual cells, yielding gene sets with shared spatial expression patterns (Methods). Cluster membership was then compared across conditions to identify genes that consistently co-clustered differently between WT and 5xFAD while showing reproducible co-clustering across biological replicates. This analysis revealed multiple condition-specific gene clusters across inhibitory neurons, excitatory neurons, and activated microglia (Table S10). In excitatory neurons, a cluster composed of *Mfap4*, *Snca*, and *Unc13a* showed coordinated spatial organization specific to the disease condition (Fig. 4B). In inhibitory neurons, *Ctsd* and *Pvalb* displayed coordinated spatial programs that differ between conditions, as did *Bsn* and *Plp1* (Fig. 4C, Fig. S5). Activated microglia also exhibited condition-specific patterning, with *Irg1*, *Neurod6*, and *Snap25* forming a gene cluster that consistently co-clustered differently in 5xFAD than in WT (Fig. S5). Together, these results suggest that early-stage 5xFAD hippocampus exhibits cell-type-specific reorganization of spatial gene neighborhoods at the single-cell level, revealing coordinated spatial programs that differ between conditions.

**Fig. 4.**
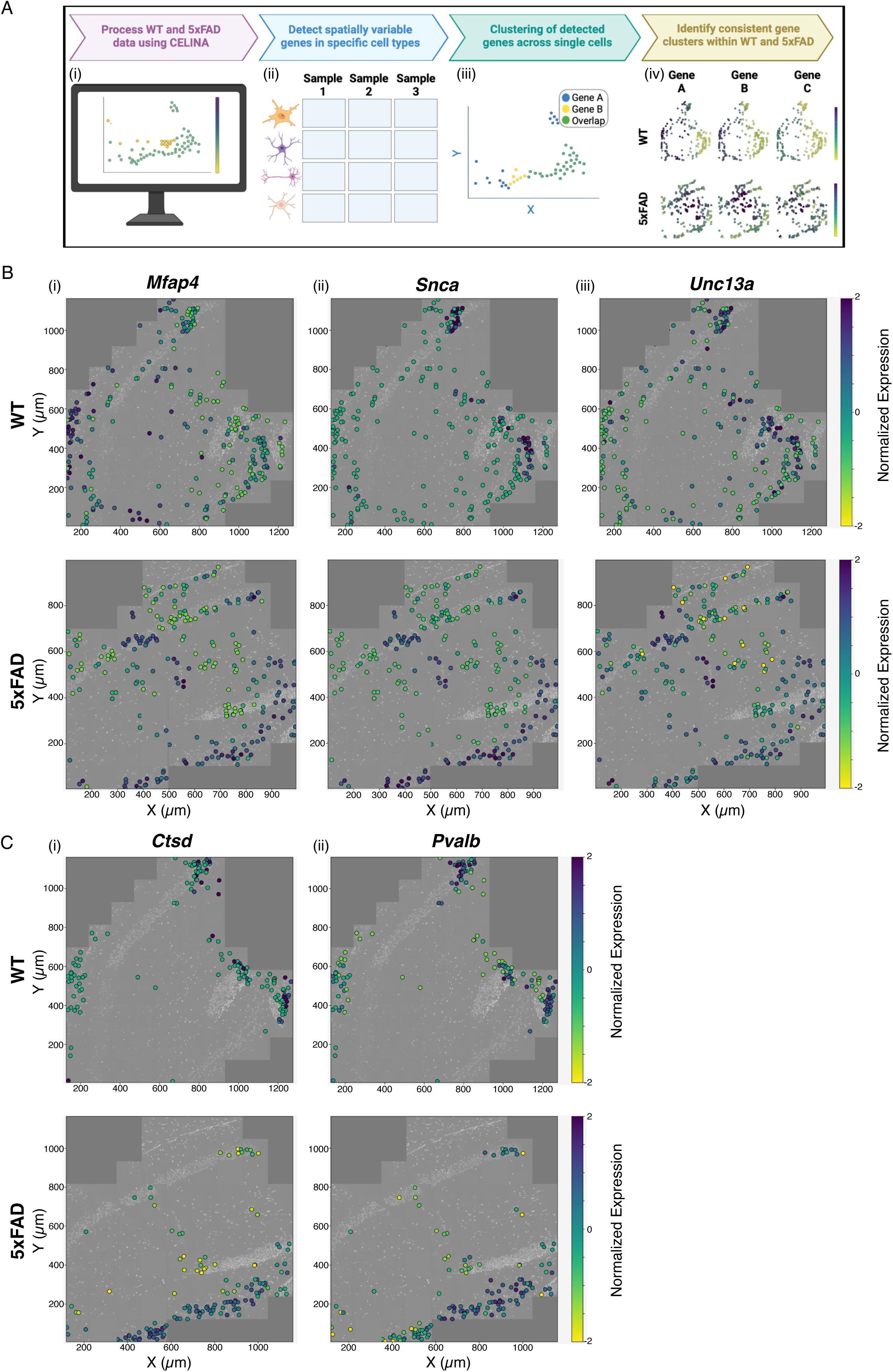
Single-cell neighborhood analysis of spatially variable gene clusters in WT and 5xFAD. **(A)** Schematic of the analysis workflow: (i) genes with spatial expression deviating from the expectation based on all genes in the same cell type were detected using Celina^43^; (ii) this was repeated for each cell type in each sample, generating a cell type × sample matrix of detected genes; (iii) within each cell type and sample, detected genes with shared expression across individual cells were clustered together; (iv) gene clusters were intersected across 5xFAD and WT samples separately to identify clusters with differential spatial patterns between conditions. **(B)** Examples in excitatory neurons: (i) *Mfap4*, (ii) *Snca*, and (iii) *Unc13a* consistently co-cluster in 5xFAD differently than in WT. **(C)** Examples in inhibitory neurons: (i) *Ctsd* and (ii) *Pvalb* show condition-specific clustering. Plots display DAPI-stained nuclei (gray) and per-cell normalized expression levels (yellow-purple scale; log-transformed and z-scaled). Top row: WT; bottom row: 5xFAD.

### Spatial RNA velocity reveals early cell type-, region-, and interaction-specific state changes

To assess whether early alterations in cellular states accompany spatial transcriptomic changes in 5xFAD hippocampus, we performed spatial RNA velocity analysis^44^ (Fig. 5). Leveraging the super-resolution of ExSeq^27^, we unambiguously distinguished nuclear and cytoplasmic compartments in each cell, a prerequisite for accurate spatial RNA velocity estimation^45^. Based on transcript abundances from these compartments, we predicted future cell states and projected them into PCA space, enabling quantification of both the magnitude and phase of cell state changes (Methods, Fig. S6, Table S11).

**Fig. 5.**
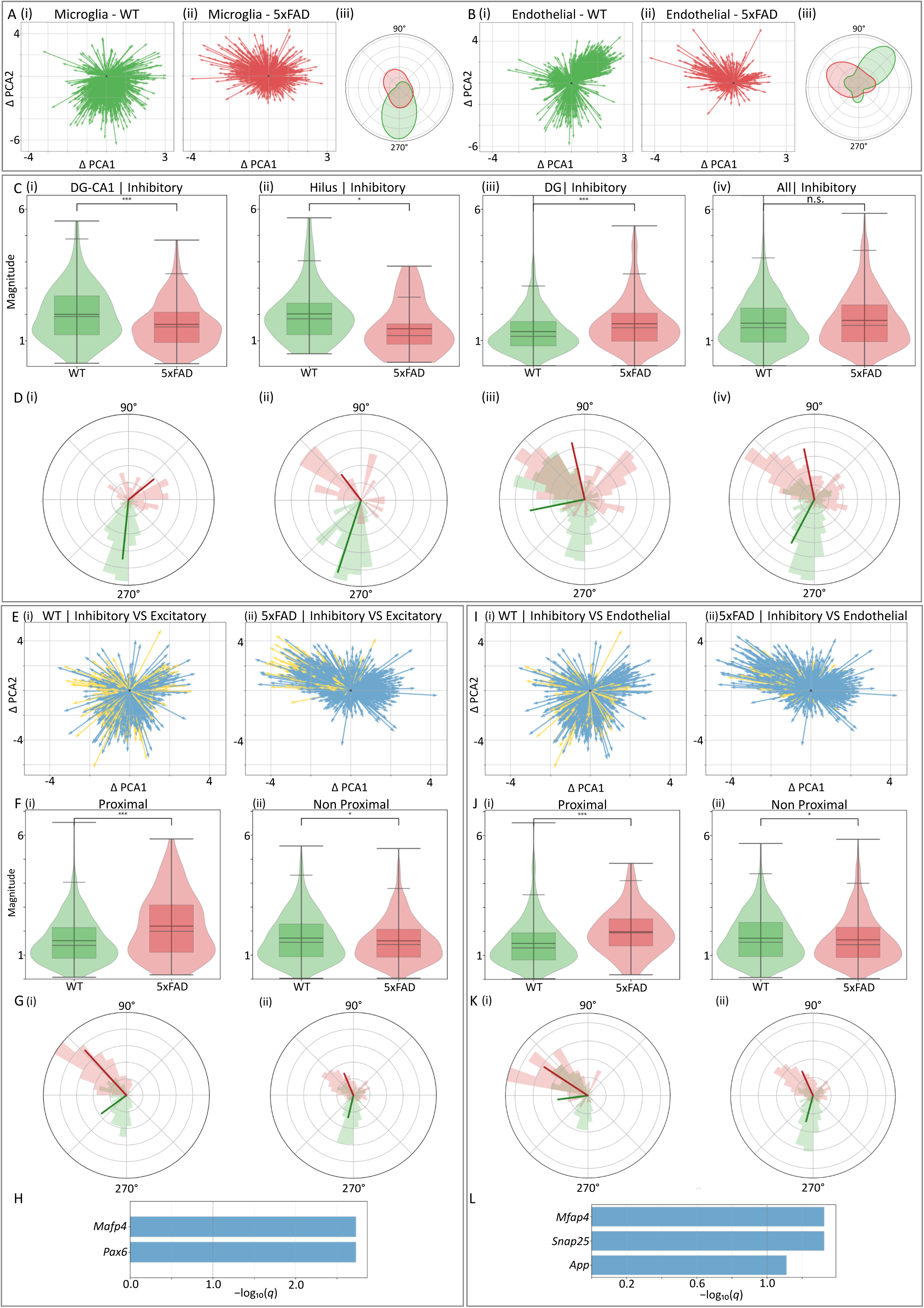
Global and region-specific cell-state differences, and their association with cell-cell proximity, between 5xFAD and control. Global hippocampus and region-specific differences in cell states between 5xFAD and control **(A-D)**. The magnitude and phase of cell state changes were assessed by spatial RNA velocity analysis for each cell type and compared between 5xFAD and control. **(A)** Microglia: quiver plots in PCA space showing state displacements for control (green, i) and 5xFAD (red, ii). Each arrow represents a single cell’s displacement in PCA space, originating from (0,0). (iii) the phase of cell state changes as polar KDE plots. **(B)** Same as (A), for endothelial cells. **(C-D)** Region-specific differences in cell states between 5xFAD and control in inhibitory neurons. **(C)** Box-violin plot comparison of the magnitude of cell state changes between 5xFAD and control within DG-CA1 (i), Hilus (ii), DG (iii) and across all hippocampal regions (iv). While differences were not significant when combining all regions, they reached significance within the specific examined regions (\**P* < 0.05;\*\**P* < 0.01;\*\*\**P* < 0.001; Mann-Whitney U [MWU] test). **(D)** Phase comparison of cell state changes using polar histograms for control (green) and 5xFAD (red) in DG-CA1 (i), Hilus (ii), DG (iii) and across all regions (iv); radial spokes indicate mean directions. Phase differences are significant in all panels displaying phase (*P* < 5 × 10^−4^, permutation analysis). Cell-cell proximity corresponds with changes in cell states between 5xFAD and control **(E-L)**. **(E)** Inhibitory neurons: changes in cell state in 5xFAD and control as a function of physical proximity to excitatory neurons. Distance-stratified quiver plots in PCA space are shown for control (i) and 5xFAD (ii); yellow = inhibitory neurons proximal (≤1 µm) to excitatory neurons; blue = inhibitory neurons non-proximal (>1 µm). **(F)** Box-violin plots comparing the magnitude of cell state changes between 5xFAD (red) and control (green) within inhibitory neurons proximal to excitatory neurons (i) and non-proximal (ii). Significant increases and decreases are observed, respectively (\**P* < 0.05; \*\**P* < 0.01; \*\*\**P* < 0.001; MWU test). **(G)** Phase comparison of cell state changes using polar histograms for control (green) and 5xFAD (red) within inhibitory neurons proximal (i) and non-proximal (ii) to excitatory neurons. Significance was assessed by a permutation test. **(H)** Genes in inhibitory neurons whose RNA velocities significantly correlated with distance to excitatory neurons (Pearson’s correlation, permutation *P*, BH-adjusted *q* values), potentially contributing to the observed cell state differences. **(I-L)** Same as (E-H), but for inhibitory neurons and their proximity to endothelial cells. Phase differences are significant in all panels displaying phase (*P* < 1 × 10^−3^, permutation analysis).

At the cell type level, RNA velocity analysis revealed clear differences in predicted future cell states between 5xFAD and controls (Fig. S7A-D), particularly in microglia (Fig. 5A) and endothelial cells (Fig. 5B). These alterations reflected both changes in the amplitude of cell state displacements and shifts in their directional phase (Fig. 5A-B). Region-specific analyses revealed additional layers of granularity that were not apparent when examining the hippocampus as a whole (Methods, Table S12-S13). Inhibitory neurons in DG-CA1 and Hilus exhibited significantly smaller velocity magnitudes in 5xFAD relative to controls, despite no differences when all hippocampal regions were pooled (Fig. 5C). Polar histograms further revealed phase shifts in DG-CA and Hilus inhibitory neurons (Fig. 5D). Comparable region-specific effects were observed in microglia within DG-CA1 and excitatory neurons within SO (Fig. S7E-J). Gene-level analysis identified transcripts with altered RNA velocities, including *Tyrobp* in microglia and *Psen2* in excitatory neurons, which may contribute to the observed cell state differences (Fig. S7E-J). These findings indicate that early transcriptomic dynamics in 5xFAD mice are not uniform across the hippocampus but instead localized to distinct subregions and cell types.

We next examined whether local cell-cell proximity influenced RNA velocity-derived cell state changes (Fig. 5, Tables S13-S15). Inhibitory neurons with cell bodies in likely direct soma-soma contact with the cell bodies of excitatory neurons (approximately less than 1 µm; Fig. S6L and Methods) exhibited significantly greater magnitudes of cell state change in 5xFAD compared with controls, whereas inhibitory neurons with cell bodies not in direct soma-soma contact with the cell bodies of excitatory neurons (>1 µm) showed reduced magnitudes (Fig. 5E-F). Phase analyses using polar histograms further revealed significant shifts in the direction of cell state transitions between 5xFAD and control for both proximal and non-proximal inhibitory neurons (Fig. 5G). At the gene level, RNA velocities of *Mfap4* and *Pax6* correlated with distance to excitatory neurons (Fig. 5H), indicating that proximity to excitatory cells can drive gene-specific velocity alterations that may contribute to the observed cell state changes. A similar pattern was observed for inhibitory neurons in relation to endothelial cells: proximal inhibitory neurons displayed increased magnitudes and altered phases of cell state change in 5xFAD compared with controls, accompanied by proximity-associated gene velocity signatures (Fig. 5I-L and additional analysis in Fig. S7K). Together, these results demonstrate that spatial RNA velocity uncovers early transcriptomic state changes in 5xFAD hippocampus that are cell type-specific, region-specific, and shaped by local cell-cell interactions.

## Discussion

Alzheimer’s disease involves progressive synaptic and neuronal loss, but the molecular alterations that precede overt pathology remain unclear. Bulk transcriptomic studies suggest that dysregulation begins weeks before plaque deposition, yet the spatial dimension of these early changes has not been systematically explored. To address this, we applied ExSeq to hippocampi of 4-week-old 5xFAD mice and wild-type littermates, a stage when hippocampal morphology is preserved and plaques are absent, to test whether subtle spatial alterations arise before classical hallmarks of disease. We detected tens of genes with altered expression in different regions of the hippocampus (Table S7). Moreover, our results show that early transcriptomic alterations in 5xFAD extend beyond changes in expression to include spatial reorganization of RNA, with 23 genes displaying altered localization despite unchanged abundance. Single-cell neighborhood analysis further uncovered gene clusters co-regulated in 5xFAD within inhibitory and excitatory neurons, highlighting condition-specific local programs. Spatial RNA velocity extended these findings by capturing early, dynamic changes in cell states, which were cell-type-and region-specific, and strongly influenced by cell-cell proximity. Of particular interest, we find that inhibitory neurons exhibit cell state changes in 5xFAD compared with controls as a function of cell-cell proximity, in line with previous reports of inhibitory neuron alterations in 5xFAD mice at advanced disease stages^46^.

### Presynaptic vesicle priming axis and potential early hyperexcitability

Our early-stage spatial analysis revealed a coordinated upregulation of presynaptic priming genes in the 5xFAD hippocampus. *Rims1* and *Unc13a (Munc13-1)* were broadly elevated across multiple regions, with *Unc13a* also showing altered spatial distribution. *Rph3a* (*Rabphilin-3A*) was increased in SLM and DG-CA1. Since RIM proteins activate Munc13 to drive vesicle docking and priming, and RPH3A promotes SNARE assembly, these changes point to a compensatory enhancement of presynaptic priming^47–51^. Such alterations may favor early hyperexcitability, consistent with reports of increased neuronal activity in early AD^52^. *Stxbp5 (tomosyn)* was found in our analysis to be elevated in 5xFAD compared with WT across several hippocampal regions, including CA1, CA3, DG-CA1, and SO, with additional localization differences in SLM. Tomosyn inhibits SNARE-complex formation by sequestering syntaxin-1, thereby reducing release probability^53^. This might represent an early stage inhibition of synaptic transmission in 5xFAD mice that might be compensated by an upregulation of proteins that support vesicle exocytosis like RIM1 and Munc13-1. This broad upregulation, together with its localization differences, may represent a homeostatic mechanism that partly counteracts the upregulated priming activity observed in 5xFAD.

### Region-specific and spatial reorganization of synaptic proteins

*Bsn* (*Bassoon*), an active zone protein^54–57^, was higher in WT than 5xFAD in CA1 and DG-CA1, but elevated in 5xFAD in CA3 and SLM, with additional localization differences in WT DG-CA1. As Bassoon helps organize vesicle pools and align release machinery with Ca^2+^ entry, these opposite trends in its mRNA spatial distribution point to region-specific remodeling of release sites, though mRNA changes may not directly reflect protein levels. *Snap25*, encoding for SNAP25 which is involved in SNARE complex formation and vesicle fusion, was found in our analysis to be elevated in WT compared with 5xFAD in SLM and DG-CA1, yet showed localization differences in 5xFAD across SLM, DG-CA1, and SO. The combination of higher overall levels in WT but spatial reorganization in 5xFAD may suggest circuit-selective remodeling rather than uniform loss. These spatial reorganizations of synaptic transcripts, together with region-specific changes, highlight the need for future studies to investigate circuit- and even synapse-specific alterations as potential early biomarkers of AD.

### APOE and α-synuclein: potential contributors to presynaptic stress

*Apoe* was found in our analysis to be elevated in 5xFAD compared with WT in CA1 and CA3, with additional localization differences in SLM and DG-CA1. APOE^58^, and especially ApoE4, is a major risk factor for AD and is known to influence Aβ aggregation, clearance, and microglial metabolism, suggesting that these changes may contribute to early vulnerability. *Snca (α-synuclein)* was also elevated in 5xFAD and showed localization differences across CA1, CA3, SLM, and DG-CA1. Given that α-synuclein co-pathology can exacerbate neurodegenerative outcomes^59^, these clustered increases may indicate early presynaptic stress hubs.

### Complement and microglia: early indications of synaptic pruning

Complement tagging and microglial activation are thought to act in concert to drive synaptic pruning^60^. *C1qa*, a gene that codes for a key component of the complement system, was found in our analysis to be elevated in 5xFAD compared with WT across CA1, CA3, DG, and SO, with additional localization differences in SLM and DG-CA1. *C1qc* also showed localization differences in SLM of 5xFAD. We next inspected our dataset for genes associated with disease-associated microglia. *Trem2* was elevated in 5xFAD in CA1, CA3, SLM, and SO, and in DG-CA1 it showed both higher expression and localization differences. *Tyrobp* was elevated in SO, with additional localization differences in SLM. Thus, *Trem2, Tyrobp,* and *C1qa* all showed elevated expression with localization differences in 5xFAD, consistent with early activation of microglial pathways that are implicated in synaptic pruning. Other genes linked to disease-associated microglia included *Laptm5* (elevated in CA1, SLM, DG-CA1, SO) and *Cst7* (localization differences in CA1). Together, these findings suggest that spatial hotspots in SLM and DG-CA1 may reflect early microglial activity associated with complement-mediated synaptic pruning.

### Oxidative and hypoxic stress signatures

Additional alterations pointed to localized redox imbalance and hypoxia-responsive signaling. *Nos1*, which codes for neuronal nitric oxide synthase, was elevated in CA1, SLM, and DG-CA1 of 5xFAD, supporting increased nitrosative stress. *Hif1a* displayed localization differences in SLM, consistent with hypoxia-driven reprogramming, while *Apod* was elevated in CA1 with additional localization differences in SLM and DG-CA1, in line with its role in antioxidant defense. In addition, response to reactive oxygen species is functionally enriched among the 23 genes exhibiting altered localization without changes in abundance in the 5xFAD hippocampus (Table S17). Together, these findings suggest that redox and hypoxic stress are spatially organized early in disease progression.

### Mitochondrial dysfunction

We also detected signs of mitochondrial stress, particularly affecting protein import. *Tomm20* was elevated in CA1 with localization differences in SLM and DG-CA1. As a receptor of the outer-membrane TOM import complex^61^, TOM20 alterations may indicate regional disruption of mitochondrial protein import at early stages.

Together, these results establish that subtle, spatially organized molecular disruptions are already present in the hippocampus before plaque formation, providing new insights into the earliest stages of AD progression. The full super-resolution ExSeq dataset, provided together with a custom graphical user interface for 3D visualization, offers a resource that will enable additional interrogation of early spatial mislocalization at the nanoscale, including transcript positioning relative to nuclei and to regional hippocampal morphology. These findings also provide input for future studies that can focus on synapse-specific and pathway-specific investigation of early and late changes in AD.

### Limitations of the study

Despite the ability of ExSeq to reveal early, spatially resolved RNA localization and cell state dynamics at single-cell resolution, several limitations should be considered. First, spatial RNA analysis does not directly capture functional consequences. Given the alterations observed in synaptic and presynaptic genes, future studies integrating RNA localization with electrophysiological measurements, and extended to larger cohorts, will be critical to link spatial transcriptomic changes to neuronal excitability and circuit function. Second, this study focused on a targeted panel of 101 genes, limiting transcriptome-wide discovery. Extending ExSeq to larger gene panels may uncover additional early spatial programs. Third, our analysis was restricted to the hippocampus at a single early time point. Since AD affects distributed brain networks over time, examining additional brain regions and disease stages will be necessary to assess the progression of the observed spatial alterations. Finally, as the 5xFAD model represents familial AD, caution is warranted when extrapolating these findings to sporadic AD.

## Resource availability

### Data and code availability

The full ExSeq dataset, including interactive data explorer, is available via Zenodo (https://doi.org/10.5281/zenodo.17568649). This repository also includes all scripts used to analyze this dataset.

## Acknowledgments

This study was supported by Brightfocus Foundation grant 929965 (U. A. and S.A.). The Israel Science Foundation (ISF) grant 2958/21 (S.A.). The Ministry of Science and Technology (MoST) grants 0005965 and 3-17970 (S.A.). European Research Council (ERC) grant 101117324 (S.A.). Israel Innovation Authority’s OrganoSpheres Consortium (S.A.). The Ministry of Innovation, Science & Technology, Israel, 1001576154 (U. A.), Israel Science Foundation, ISF grant: 2141/20 (U. A.), NIH grant 1R21AG074846-01A1 (U. A.), the Michael J. Fox Foundation, MJFF-022407 (U. A.), TEVA Pharmaceutical Industries (U. A.), and The Aufzien Family Center for the Prevention and Treatment of Parkinson’s Disease at Tel Aviv University (U. A).

